# Conductance mechanisms of rapidly desensitizing cation channelrhodopsins from cryptophyte algae

**DOI:** 10.1101/2020.03.20.001099

**Authors:** Oleg A. Sineshchekov, Elena G. Govorunova, Hai Li, Yumei Wang, Michael Melkonian, Gane K.-S. Wong, Leonid S. Brown, John L. Spudich

## Abstract

Channelrhodopsins guide algal phototaxis and are widely used as optogenetic probes for control of membrane potential with light. “Bacteriorhodopsin-like” cation channelrhodopsins (BCCRs) from cryptophytes differ in primary structure from other CCRs, lacking usual residues important for their cation conductance. Instead, BCCR sequences match more closely those of rhodopsin proton pumps, containing residues responsible for critical proton transfer reactions. We report 19 new BCCRs, which, together with the earlier 6 known members of this family, form three branches (subfamilies) of a phylogenetic tree. Here we show that the conductance mechanisms in two subfamilies differ with respect to involvement of the homolog of the proton donor in rhodopsin pumps. Two BCCRs from the genus *Rhodomonas* generate photocurrents that rapidly desensitize under continuous illumination. Using a combination of patch clamp electrophysiology, absorption and Raman spectroscopy, and flash photolysis, we found that the desensitization is due to rapid accumulation of a long-lived nonconducting intermediate of the photocycle with unusually blue-shifted absorption with a maximum at 330 nm. These observations reveal diversity within the BCCR family and contribute to deeper understanding of their independently evolved cation channel function.

**IMPORTANCE:** Cation channelrhodopsins, light-gated channels from flagellate green algae, are extensively used as optogenetic photoactivators of neurons in research and recently have progressed to clinical trials for vision restoration. However, the molecular mechanisms of their photoactivation remain poorly understood. We recently identified cryptophyte cation channelrhodopsins, structurally different from those of green algae, which have separately evolved to converge on light-gated cation conductance. This study reveals diversity within this new protein family and describes a subclade with unusually rapid desensitization that results in short transient photocurrents in continuous light. Such transient currents have not been observed in the green algae channelrhodopsins and are potentially useful in optogenetic protocols. Kinetic UV-vis spectroscopy and photoelectrophysiology reveal the desensitization is caused by rapid accumulation of a non-conductive photointermediate in the photochemical reaction cycle. The absorption maximum of the intermediate is 330 nm, the shortest wavelength reported in any rhodopsin, indicating a novel chromophore structure.

## INTRODUCTION

Channelrhodopsins are light-gated channels first discovered in green (chlorophyte) flagellate algae, in which they serve as photoreceptors mediating phototaxis by depolarization of the cell membrane (1–3). Currently, channelrhodopsins are widely used for control of neurons and other excitable cells with light (“optogenetics”) (4) for research and also in clinical trials to restore vision to the blind (5). Channelrhodopsins from chlorophyte algae conduct cations and are therefore referred to as cation channelrhodopsins (CCRs). Anion-conducting channelrhodopsins (ACRs) have been found in the phylogenetically distant cryptophyte algae (6) and a second family more recently in environmental DNA samples of unidentified origin (7). These three channelrhodopsin families share ~50% of overall sequence homology, including several key residues shown to be required for their channel activity.

However, cryptophyte genomes also encode a family of microbial rhodopsins that show a higher sequence homology to haloarchaeal proton-pumping rhodopsins than to any known channelrhodopsins, and yet exhibit cation channel activity, apparently a product of convergent evolution (8–10). In particular, these proteins contain homologs of the two carboxylate residues that serve as the Schiff base proton acceptor and donor in *Halobacterium salinarum* bacteriorhodopsin (Asp85 and Asp96, respectively), which together with the Thr89 homolog form the “DTD” motif characteristic of proton pumps. In contrast, in all other known channelrhodopsins one or both of these positions are occupied by non-carboxylate residues.

Earlier we have shown that channel activity in CCRs 1 and 2 from the cryptophyte alga *Guillardia theta* (*Gt*CCR1 and *Gt*CCR2) is mechanistically distinct from that in chlorophyte CCRs (9). According to our model, channel opening in these proteins requires deprotonation of the Asp96 homolog and occurs >10-fold faster than reprotonation of the retinylidene Schiff base. The latter is achieved by return of the proton from the earlier protonated acceptor, thus preventing vectorial proton translocation across the membrane. To emphasize their distinction from other known CCRs, we named these proteins “<u>b</u>acteriorhodopsin-like <u>c</u>ation <u>c</u>hannel<u>r</u>hodopsins” (BCCRs) (9).

Besides their fundamental importance as independently evolved light-gated cation channels, BCCRs have attracted attention as optogenetic tools, because some of them exhibit more red-shifted absorption enabling use of deeper penetrating long wavelength light and a higher Na^+^/H^+^ permeability ratio favorable for neuron depolarization with minimal acidification (8,11) compared to blue-light activated channelrhodopsin 2 from *Chlamydomonas reinhardtii (Cr*ChR2), the molecule that so far has been most popular in optogenetic studies (12). Recently ChRmine, a BCCR, has been used successfully to activate mouse neocortical neurons with orange light (13).

Here we describe 19 BCCRs from 9 cryptophyte species, the protein sequences of which form two separate branches (subfamilies) of the phylogenetic tree (Fig. 1A) in addition to that comprising the previously characterized *Gt*CCR1 and *Gt*CCR2. Two of these proteins derived from two *Rhodomonas* species utilize a different activation mechanism and exhibit rapid desensitization of photocurrent under continuous illumination. We show that the photochemical cycle of these channelrhodopsins involves accumulation of an extremely short-wavelength-absorbing and long-living intermediate responsible for fast inactivation of their photocurrents. These observations reveal diversity within the BCCR family and contribute to deeper understanding of their cation channel function independently evolved from chlorophyte CCRs.

**Figure 1.**
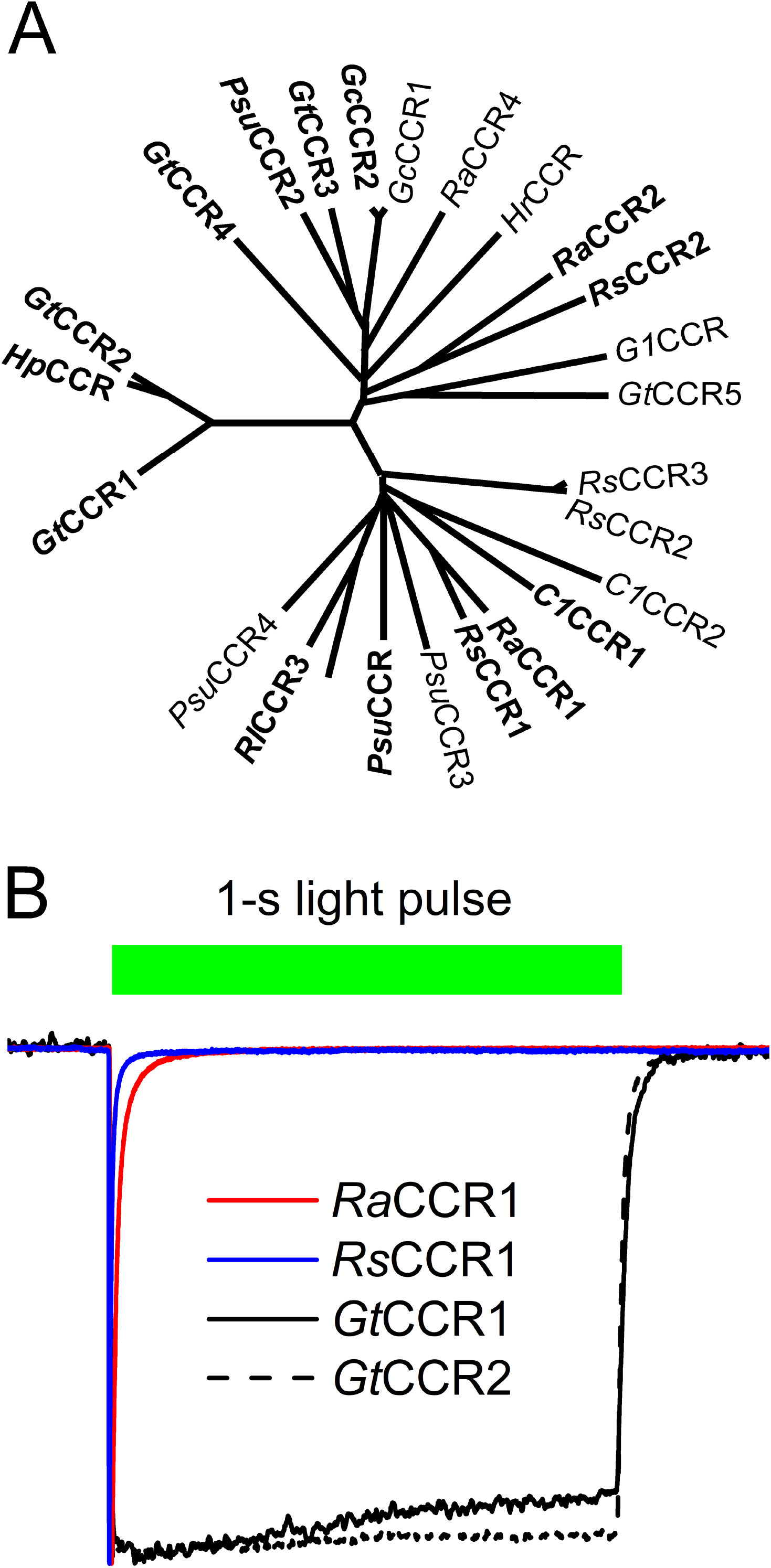
(A) A phylogenetic tree of BCCR transmembrane domains. Bold font shows the proteins that generated photocurrents upon expression in mammalian cells. (B) Normalized photocurrent traces from *Ra*CCR1 and *Rs*CCR1 (colored lines) recorded at −60 mV in response to a light pulse, duration of which is shown as the bar on top. Traces from previously characterized *Gt*CCR1 and *Gt*CCR2 (black lines) are shown for comparison.

## RESULTS

### Identification and electrophysiological screening of BCCR homologs

Using probabilistic inference methods based on profile hidden Markov models (14) built on previously known BCCR sequences from *G. theta*, we identified 19 new BCCR homologs from nine marine cryptophyte strains included in the ongoing algal transcriptome sequencing projects (15,16). The majority were cold-water species (from the Arctic or Antarctic), but *Rhodomonas lens* was from the Gulf of Mexico and *R. salina*, from Milford, Connecticut. The previously unclassified strain CCMP 2293 has recently been allocated to the new genus *Baffinella* (as *B. frigidus*) (17).

Table 1 lists GenBank accession numbers, source organisms, transcript names, and abbreviated protein names of the BCCR homologs identified in this study. In the abbreviated protein names, the first two or three letters stand for the beginning letters of the genus and species name. One of the sequences derived from *Rhodomonas lens* (*Rl*CCR1) exactly matched the sequence recently reported under the name “ChRmine” and attributed to the marine ciliate *Tiarina fusus* (13). As *T. fusus* culture used for RNA isolation was fed on *R. lens* (biosample SAMN02740485), the presence of this sequence in the *T. fusus* transriptome can be explained by insufficient starvation of the organisms prior to RNA extraction.

**Table 1.**
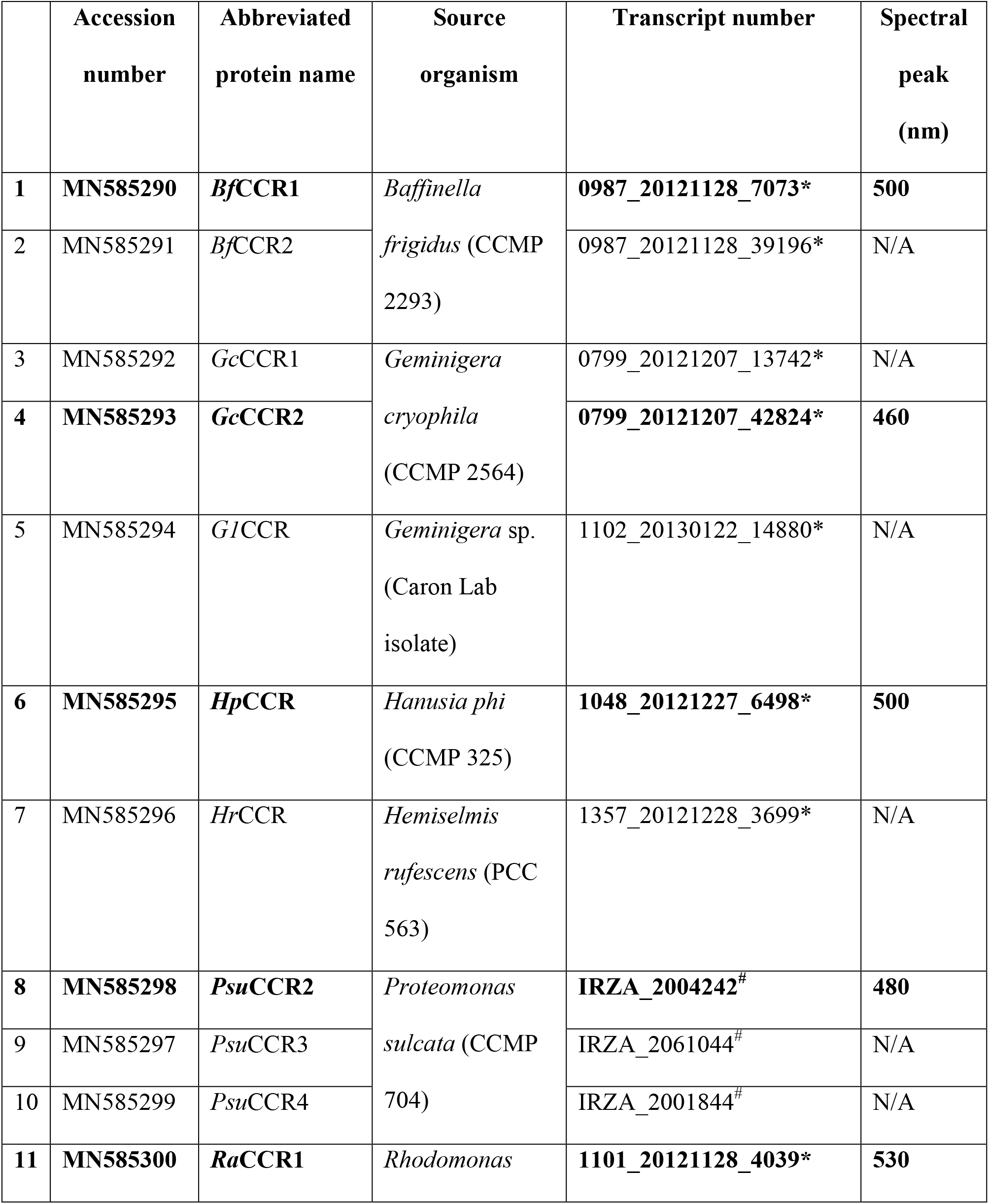
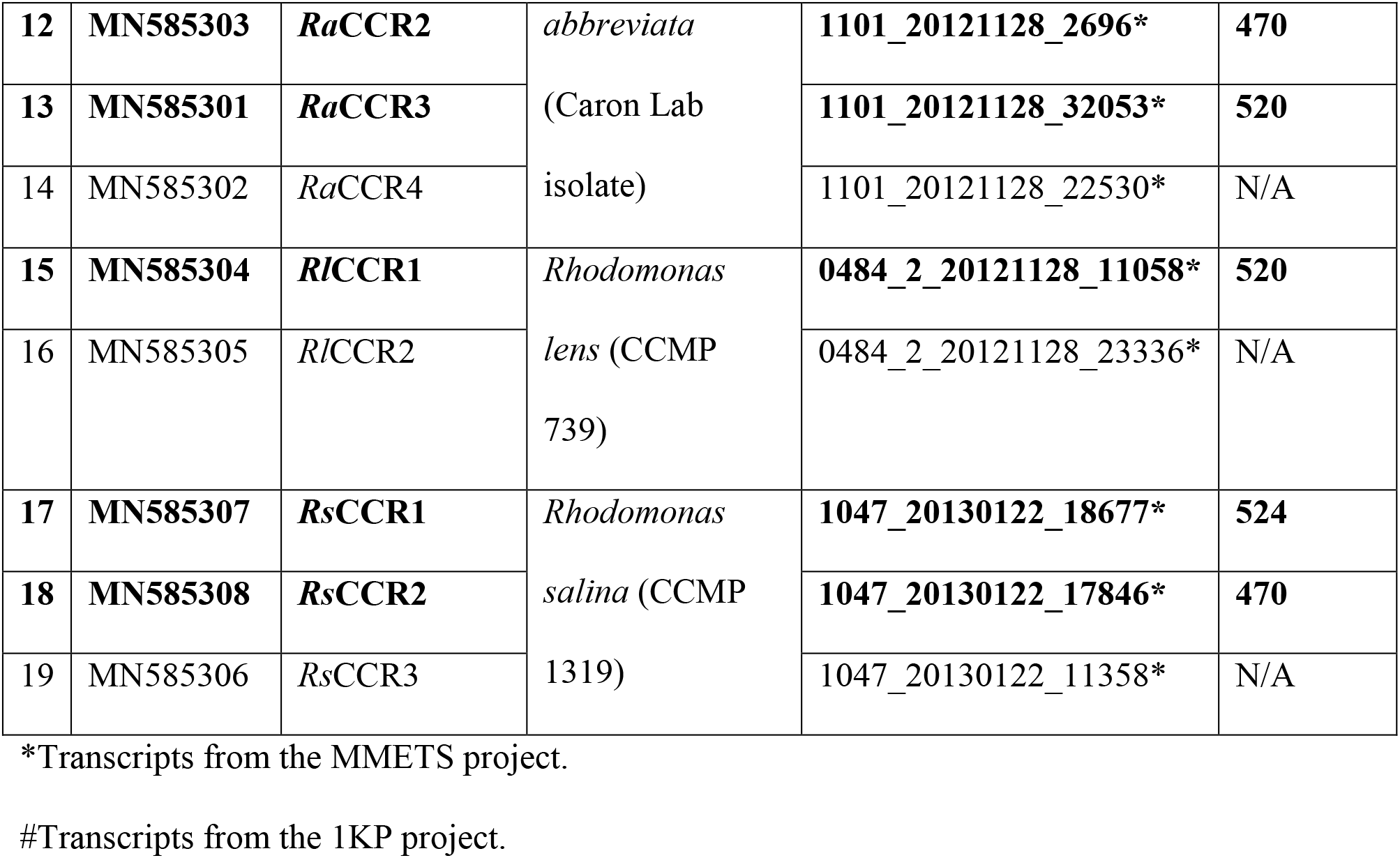
A list of BCCR homologs tested in this study (functional homologs are in bold font).

Fig. S1 shows protein alignment of the opsin domains of BCCRs identified in this study. Asp85 and Thr89 (bacteriorhodopsin numbering) are conserved in all sequences, whereas Asp96 is replaced with Glu and Thr in *Bf*CCR2 and *G1*CCR, respectively. However, neither of these sequences were electrogenic upon expression in mammalian cells (see below), and therefore the functional importance of these substitutions could not be assessed. In many BCCR homologs the position of bacteriorhodopsin’s Arg82 is occupied by other residues (Lys, Ala, Pro, Gln or even Glu (in *Ra*CCR2)), which is unusual among microbial rhodopsins. Fig. 1A shows a phylogenetic tree of the transmembrane domains of so far identified BCCRs. The previously characterized *Gt*CCR1 and *Gt*CCR2, together with *Hp*CCR, a closely related sequence from *Hanusia phi*, form a separate branch of this tree.

We synthesized human codon-optimized polynucleotides encoding the opsin domains of 19 newly identified BCCRs, fused them to an in-frame C-terminal EYFP (enhanced yellow fluorescent protein) tag and expressed in HEK293 (human embryonic kidney) cells. Ten of the encoded proteins generated photocurrents, the largest of which were the peak currents from *Ra*CCR1 and *Rs*CCR1 (Fig. S2A). The action spectra were determined by measuring the initial slopes of photocurrent in the linear range of the light intensity. The spectrum of *Ra*CCR1 closely matched that of *Rl*CCR1 (also called ChRmine (13)) and peaked at ~530 nm; that of *Rs*CCR1 was ~5 nm blue-shifted (Fig. S2B). The spectral maxima of other tested BCCRs are listed in Table 1. The current kinetics was very diverse in the tested BCCRs. In particular, currents recorded from *Ra*CCR1 and *Rs*CCR1 exhibited extremely rapid desensitization during continuous illumination (Fig. 1B, red and blue lines, respectively). Representative photocurrent traces from other tested BCCRs are shown in Fig. S3.

To test relative permeability for Na^+^, we partially replaced this ion in the bath with non-permeable *N*-methyl-D-glucamine (NMG^+^) and determined the reversal potentials (E_rev_) by measuring the current-voltage relationships in four BCCR variants that generated the largest photocurrents. *Ra*CCR1 and *Rs*CCR1 showed large E_rev_ shifts towards the new equilibrium potential for Na^+^ (Fig. S4A), similar to the earlier reported *Gt*CCRs (9,10). However, *Ra*CCR2 and *Rs*CCR2 showed smaller shifts, similar to that in *Cr*ChR2 (18). When we reduced the bath pH without changing its Na^+^ concentration, E_rev_ shifts were correspondingly smaller in *Ra*CCR1 and *Rs*CCR1, as compared to *Ra*CCR2 and *Rs*CCR2 (Fig. S4B), indicating a higher Na^+^/H^+^ permeability ratio of the former two CCRs, as compared to the latter. No change in E_rev_ was detected upon partial replacement of Cl^−^ in the bath with bulky aspartate, indicating that neither of the tested BCCRs conducted Cl^−^ (Fig. S4C).

### Absorption spectroscopy of RaCCR1 and RsCCR1

To gain more insight into mechanisms of their photoactivation, we expressed and detergent-purified *Ra*CCR1 and *Rs*CCR1 from the methylotrophic yeast *Pichia pastoris*. Their absorption spectra in the visible range closely matched the action spectra of photocurrents with main peaks at 530 and 524 nm, respectively (Fig. 2A). In addition, the absorption spectra of dark-adapted *Ra*CCR1 and *Rs*CCR1 showed structured absorption in the near-UV range (with peaks at ~307, 321 and 337 nm), in contrast to that of previously characterized *Gt*CCR2 purified from the same expression host.

**Figure 2.**
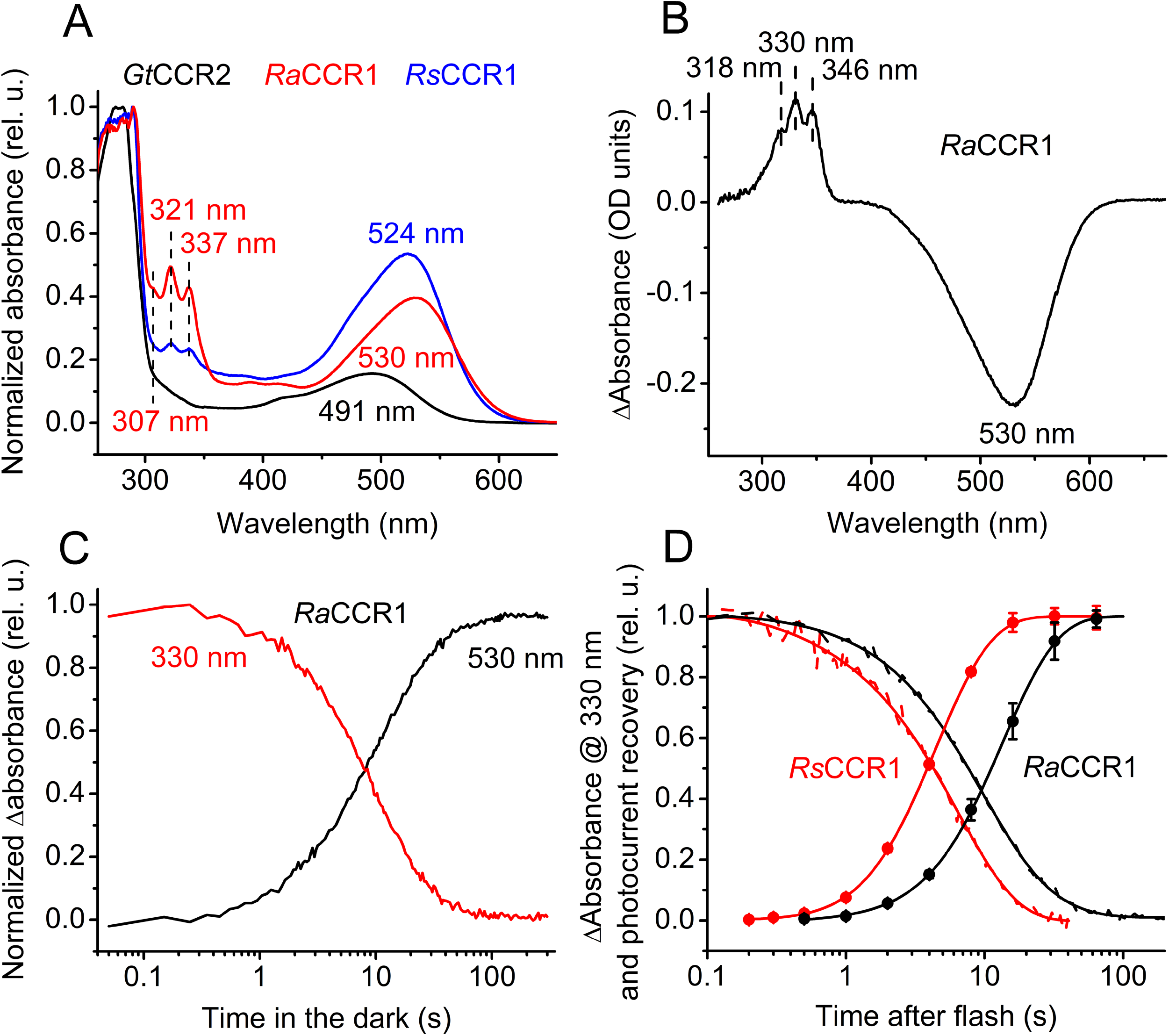
(A) The absorption spectra of dark-adapted detergent-purified proteins. (B) The difference (light minus dark) absorption spectrum of *Ra*CCR1. (C) The time course of absorption changes at 330 and 530 nm during dark incubation of illuminated *Ra*CCR1. (D) The time course of absorption changes at 330 nm in *Rs*CCR1 (that in *Ra*CCR1 from panel C is shown for comparison) and photocurrent recovery for both channelrhodopsins.

Similar UV bands had been reported in *Gt*CCR4 and tentatively attributed to impurities of the sample (10). However, incubation of *Ra*CCR1 and *Rs*CCR1 with hydroxylamine, an agent known to cleave the retinal chromophore from the bacteriorhodopsin apoprotein in a light-dependent manner (19), decreased absorption in the UV region with the difference spectrum exhibiting the same triple-peak structure characteristic of protein-bound retinal (Fig. S5A and B). In both proteins the rate of hydroxylamine bleaching of the UV bands was at least twice as fast as that of the main band (Fig. S5C and D), indicating that the UV-absorbing fractions were more accessible to hydroxylamine than the fractions absorbing in the visible range. Illumination accelerated bleaching in the visible range, as expected for retinylidene proteins, but did not influence the rate of bleaching at 321 nm (Fig. S5C and D). These results suggest that the structured UV absorbance in the dark-adapted sample is attributable to retinal binding to partially misfolded *Ra*CCR1 and *Rs*CCR1. The ratio of the UV absorption to the main peak absorption varied from 0.4 to 1.5 in different preparations, did not depend on the length of the expression construct, purification procedure, or storage conditions, and may reflect the relative amount of misfolded protein.

Continuous illumination of detergent-purified *Ra*CCR1 with visible light decreased absorption at the main band (P530) and led to formation of a product (P330) with structured absorption in the UV region with three peaks at 318, 330 and 346 nm (Fig. 2B), red-shifted from those observed in the dark. Dissipation of P330 occurred on the time scale of seconds in parallel with recovery of the unphotolyzed state P530 (Fig. 2C). The recovery of the unphotolyzed state was ~3-fold slower in *Pichia* membranes than in detergent-purified protein (Fig. S6A). A very similar product with structured UV absorption was also formed upon illumination of purified *Rs*CCR1 (Fig. S6B), with a rate of dissipation in the dark >2-fold faster than that in *Ra*CCR1 (Fig. 2D).

### Mechanism of photocurrent desensitization

As described above, photocurrents from *Ra*CCR1 and *Rs*CCR1 exhibited rapid desensitization under continuous light (Fig. 1B). Desensitization was also observed under stimulation with 6-ns laser flashes at 0.1 Hz frequency even at 10% power (Fig. S6C), which argues against its origin from a secondary photochemical process. An alternative explanation for photocurrent desensitization is the existence of a long-lived non-conductive state in the single-turnover photocycle. To determine the rate of peak current recovery, a second flash was applied after a variable time delay. The rate of restoration of the ability to generate electric current closely matched that of P330 dissipation in both purified proteins (Fig. 2D), strongly suggesting that accumulation of P330 is responsible for the rapid desensitization of photocurrents generated by *Ra*CCR1 and *Rs*CCR1.

Fig. S7A shows a series of photocurrent traces generated by *Ra*CCR1 in response to 1-s light pulses of different intensities. The peak photocurrent increased over the entire tested intensity range, whereas the degree of desensitization reached saturation two orders of magnitude earlier (Fig. S7B), and similar results were obtained with *Rs*CCR1 (Fig. S7C). These observations show that the long-lived non-conductive P330 is not in equilibrium with the unphotolyzed state of the protein.

Alkalization caused formation of a UV-absorbing species of *Ra*CCR1 with a structured spectrum closely matching that of the form obtained by illumination (Fig. 3A). The pK_a_ of this process was identical to that of decrease of absorbance at 530 nm, which showed that the UV-absorbing species was produced from P530 (Fig. 3B). The alkali-induced conversion of the unphotolyzed form absorbing at 530 nm to P330 decreased the amplitude of the photo-induced conversion. When illuminated *Ra*CCR1 samples were incubated in the dark at high pH, only a small decrease in absorbance at 330 nm was observed, as compared to neutral pH conditions (Fig. 3C). The pK_a_ of this decrease in amplitude was identical to that of conversion of P530 to the UV-absorbing species (Fig. 3D). We conclude from these observations that the same species accumulated at high pH as that obtained by illumination (i.e. P330). To the best of our knowledge, P330 is the shortest wavelength intermediate observed in the photocycle of any microbial rhodopsin. In addition to formation of P330, alkalization caused accumulation of an M-like intermediate absorbing at ~390 nm with pK_a_ ~9.0, although its concentration (assuming ~equal extinction coefficients) was 10-fold smaller than that of P330 (Fig. 3E).

**Figure 3.**
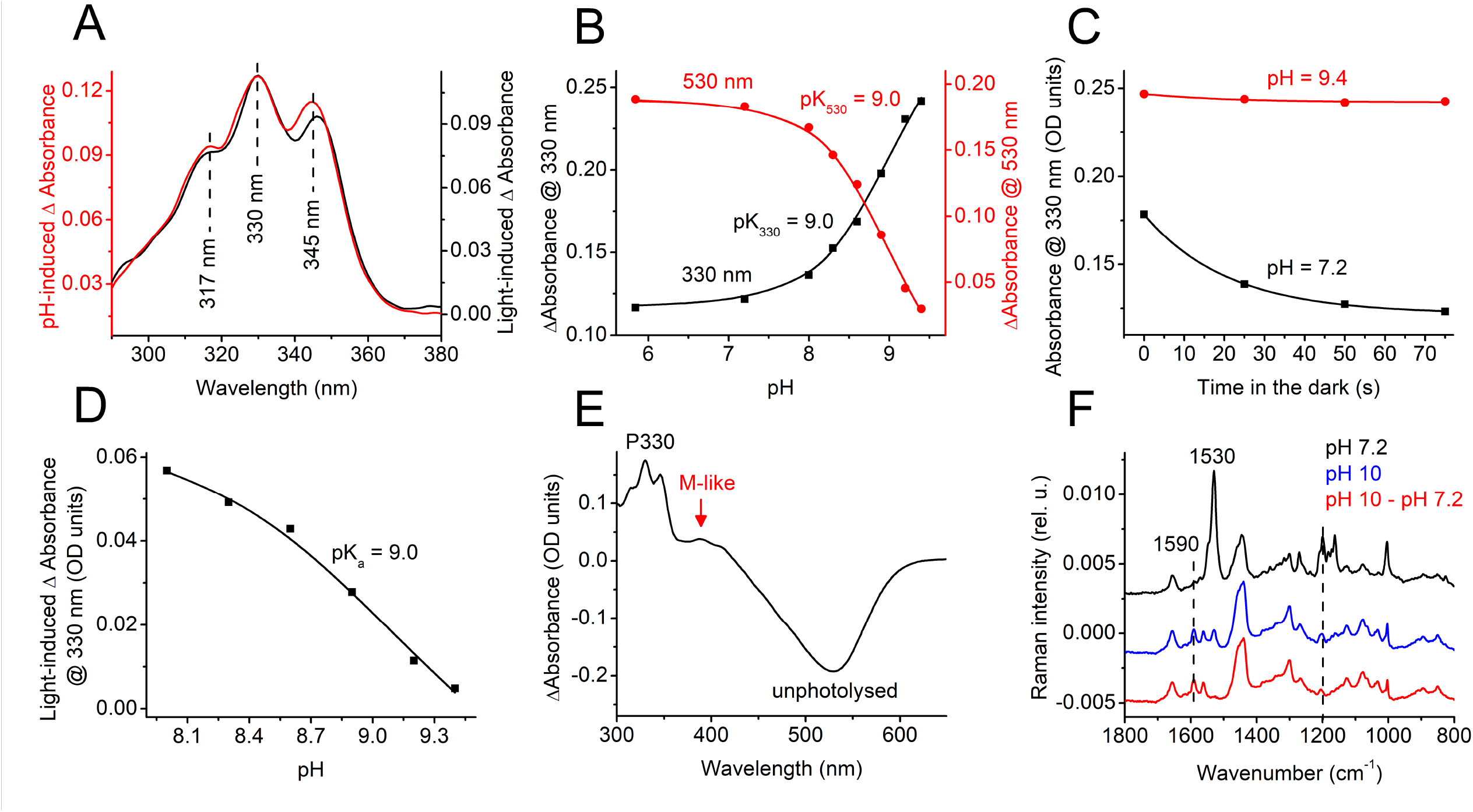
(A) The UV region of the difference absorption spectra of *Ra*CCR1 obtained upon a pH increase from 7.2 to 9.3 (red, left axis) or upon illumination (black, right axis). (B) The pH dependence of absorbance changes at 330 nm (black, left axis) and 530 nm (red, right axis). (C) Absorbance changes at 330 nm during incubation of *Ra*CCR1 in the dark at the indicated pH. (D) The pH dependence of the light-induced absorbance changes at 330 nm. (E) The difference absorption spectrum pH 10 minus pH 7.2. (F) The FT-Raman spectra measured at pH 7.2 and 10, and their difference spectrum.

At pH 10 essentially all molecules were converted from the unphotolysed form to P330. This allowed us to use FT-Raman spectroscopy to probe its chromophore structure in the dark. The Raman spectra measured at pH 7.2 and 10 and their difference spectrum are shown in Fig. 3F. The main ethylenic C=C stretch at 1530 cm^−1^, which corresponds to the main visible peak at 530 nm (20), and the fingerprint C-C stretches at 1200 and 1163 cm^−1^ showed that at pH 7.2 retinal was predominantly in an all-*trans* configuration. Upon alkalization the band at 1530 cm^−1^ was strongly reduced, and new bands appeared at 1590 cm^−1^ and 1562 cm^−1^ which presumably corresponded, respectively, to P330 and the M-like intermediate absorbing at ~390 nm. The same two bands were clearly resolved in the difference spectrum.

### Fast photochemical conversions

Fast photochemical conversions in the near UV and visible range were analyzed by flash photolysis. Fig. S8A shows a series of absorption changes in *Rs*CCR1 detected at wavelengths from 390 to 570 nm at 10-nm increments. Only negligible (less than 0.5 mOD) oppositely directed components with the time constant (τ) values ~60-100 μs were observed at the wavelengths at which maximal absorption of the red-shifted K and blue-shifted L intermediates are expected (480 and 560 nm, respectively) (Fig. S8B). Therefore we could not follow the K to L transition, which occurred on a much faster time scale. To obtain the spectral changes due to L formation, we plotted the mean absorption changes in the time window between 50 and 100 μs after the flash against wavelength (Fig. 4A). The maximum of the L intermediate in this difference spectrum was at ~460 nm.

**Figure 4.**
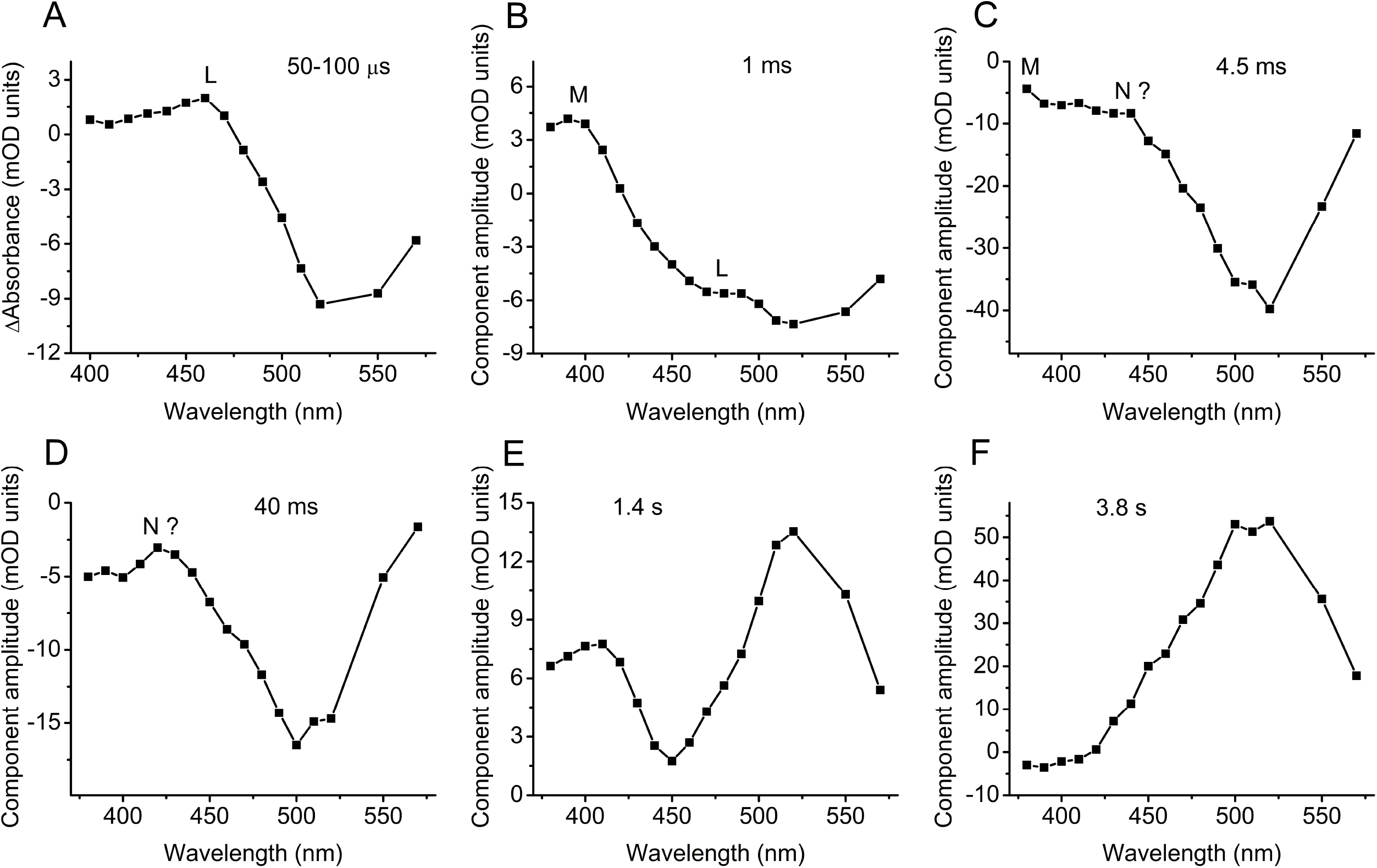
(A) Mean photoinduced absorbance changes recorded from purified *Rs*CCR1 in the 50-100 μs time window. (B-F) Spectral transitions in *Rs*CCR1 derived by global fit analysis.

The spectral characteristics of the later transitions were obtained by global fit analysis. The appearance of a typical M intermediate with the absorption maximum at ~390 nm (the positive peak in Fig. 4B) was observed within 1 ms. After that, biphasic bleaching at all visible wavelengths took place, which was obviously related to generation of P330 form. Fast bleaching with τ ~4.5 ms reflected the decay of the bulk of the initial form and may involve the appearance of a blue-absorbing (N?) intermediate (Fig. 4C), which was more obvious during the slow bleaching with τ ~40 ms (Fig. 4D). The recovery of the initial state proceeded in two steps with τ 1.4 and 3.8 s. At least the fast recovery involved depletion of a blue-absorbing intermediate (Fig. 4E). The τ of the main slow recovery component was equal to those of P330 dissipation and restoration of electrical sensitivity (Fig. 4F and Fig. 2D). Qualitatively similar phototransitions were observed in the second pigment *Ra*CCR1 with time constants of components 0.3, 6, 40, 3200 and 1140 ms (Fig. S9). In agreement with slower dissipation of the P330 intermediate and restoration of light sensitivity in this pigment as compared to *Rs*CCR1 (Fig. 2D), the recovery in the visible range was also slower, and depletion of the blue absorbing form was also observed (Fig. S9E and F). However, the 40-ms component which in *Rs*CCR1 corresponded to slow bleaching, in *Ra*CCR1 revealed fast recovery.

We recorded photocurrents in HEK cells upon 6-ns laser flash excitation at 532 nm as in flash-photolysis measurements for kinetic comparison with absorption changes in purified proteins. Channel opening and closing in *Ra*CCR1 and *Rs*CCR1 took place in the same time windows as absorption changes at the wavelengths of M-intermediate absorption in which proton transfers occur (Fig. 5A and B).

**Figure 5.**
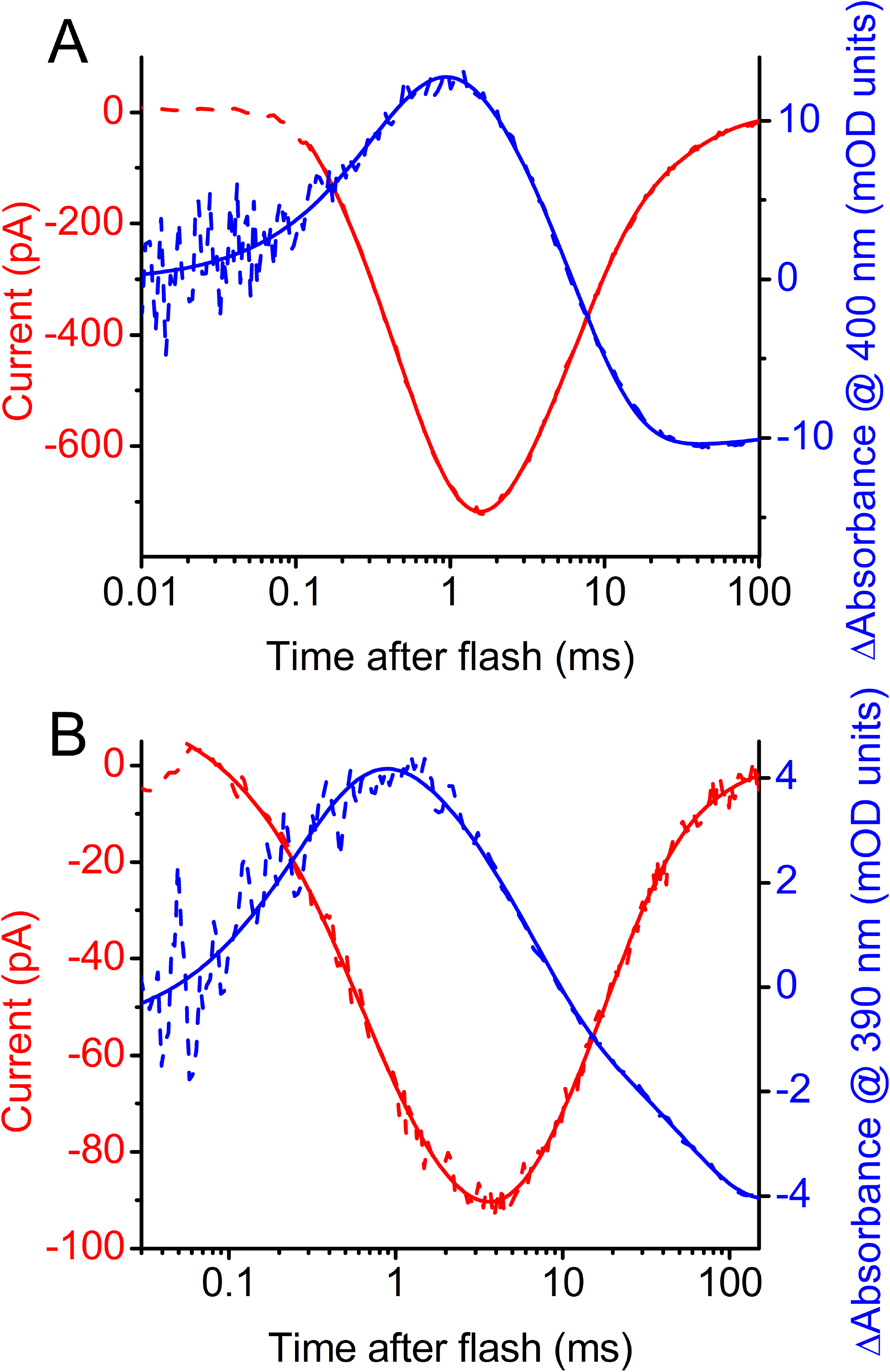
(A and B) Laser flash-evoked photocurrents at −60 mV (red) and photoinduced absorbance change (blue) recorded from *Ra*CCR1 (A) and *Rs*CCR1 (B).

In the current traces generated by *Gt*CCR1 and *Gt*CCR2, a large peak was observed in the 30-100 μs time domain prior to channel opening (9). This peak, also exhibited by some low-efficiency CCRs from green algae, reflects intramolecular transfer of the Schiff base proton to an outwardly located acceptor, integrated by the measuring system (21). This component could also be resolved in the current traces from *Ra*CCR1 and *Rs*CCR1 recorded at the voltages near the reversal potential for Na^+^, but it was ~100-fold smaller than that in *Gt*CCR1 and *Gt*CCR2 (Fig. S10).

### Mutagenesis analysis

In *Gt*CCR1 and *Gt*CCR2 we found that a neutralizing mutation of the Asp96 homolog (Asp98) completely suppressed channel activity, so that only intramolecular transfer of the Schiff base proton could be detected (9). The corresponding D125N and D128N mutations in *Ra*CCR1 and *Rs*CCR1, respectively, did not eliminate channel currents (Fig. 6A). Neutralization of the Asp85 homolog in *Ra*CCR1 (the D114N mutation) reduced expression of the construct, as judged by the tag fluorescence, and no photocurrents above the noise level could be detected. Replacement of Cys119 (corresponding to Thr90 in bacteriorhodopsin) with Ala completely abolished photocurrents in *Ra*CCR1, as did also the corresponding mutations in *Ra*CCR2, *Gt*CCR2 and *Psu*CCR. In the *Ra*CCR1_C119T mutant the photocurrent amplitude was greatly reduced (the mean peak current in response to a first light pulse of maximal intensity was 10 ± 3 pA, n = 14 cells). These tiny currents, however, exhibited only ~40% desensitization during 1-s continuous illumination (Fig. 6B), i.e., much less than in the wild type. The photocurrent decay after switching the light off was biphasic, with the slow phase on the second time scale. But the most striking difference of this mutant from the wild type was the absence of the long-living UV-absorbing form with the structured spectrum corresponding to P330 in the wild type (Fig. 6C).

**Figure 6.**
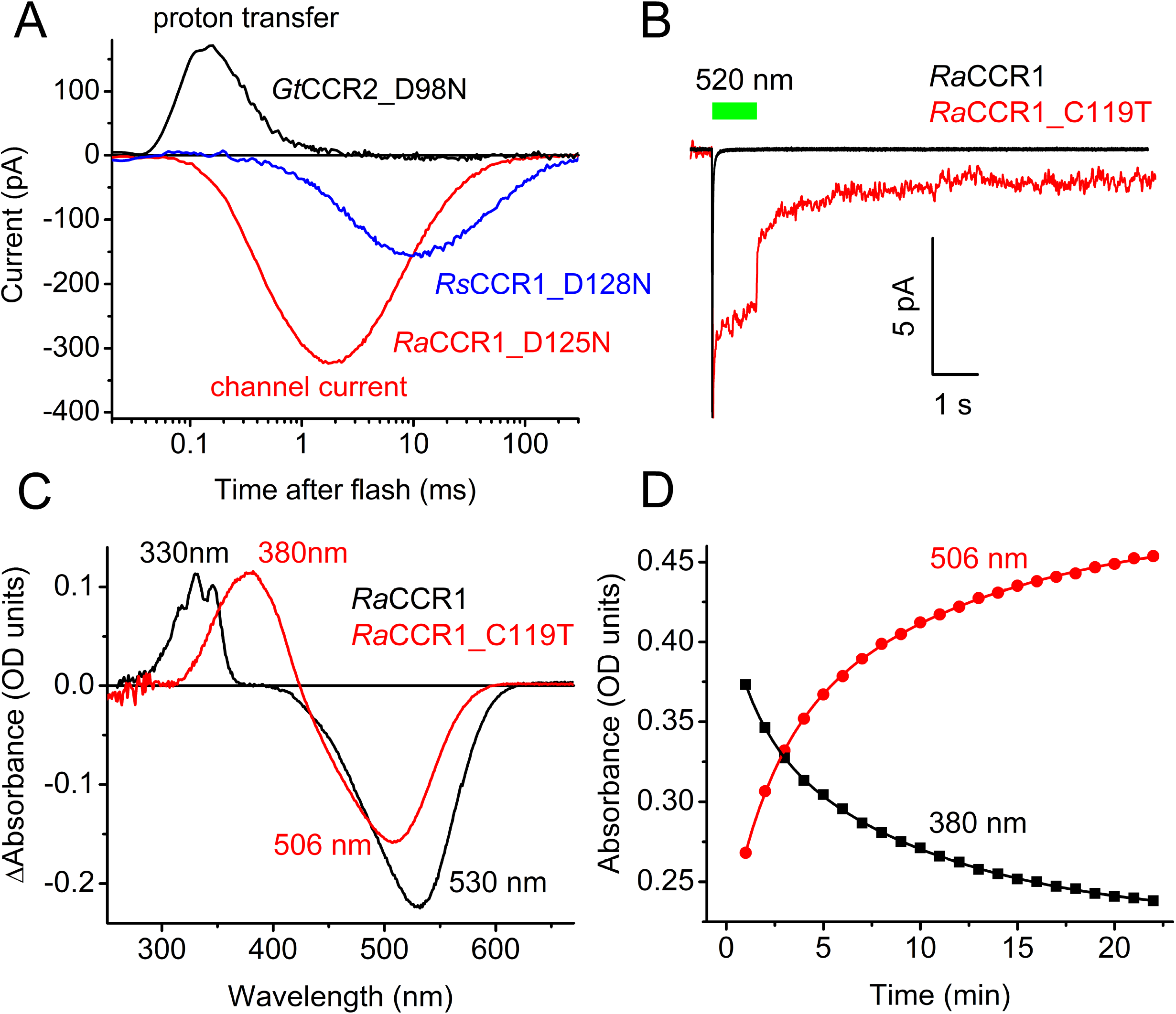
(A) Laser-evoked photocurrents recorded at −60 mV from the mutants of the Asp96 homolog in three BCCR mutants in which the homolog of Asp96 was neutralized. (B) The current trace recorded in response to 1-s light pulse from the *Ra*CCR1_C119T mutant (red line). The normalized trace from the wild type is shown in black for comparison. (C) The light minus dark absorption spectrum of purified *Ra*CCR1_C119T mutant (red). The spectrum for the wild type is shown in black for comparison. (D) Absorbance changes in purified illuminated *Ra*CCR1_C119T during incubation in the dark.

Instead, a smooth peak with the maximum at 380 nm (the M state) was produced. The dissipation of this state and the recovery of the unphotolyzed state with the peak absorbance at 506 nm were very slow (Fig. 6D).

## DISCUSSION

Our results show that BCCRs are widely spread among cryptophyte algae and form three branches (subfamilies) of a phylogenetic tree. BCCRs exhibit diverse current kinetics, spectral sensitivity and Na^+^/H^+^ permeability ratios, as has also been found in other channelrhodopsin families. Two representatives of the currently studied BCCRs differ in their mechanism of photoactivation from previously described *Gt*CCR1 and 2, which belong to a different subfamily of cryptophyte CCRs. Most notably, a particular proton transfer essential to trigger channel opening in the earlier reported subfamily is not required in the subfamily described in this study. Photocurrents by *Ra*CCR1 and *Rs*CCR1 exhibit very rapid desensitization under continuous illumination, which we show is related to the formation of a long-living UV-absorbing intermediate in their photocycles. Similar rapid photocurrent desensitization was observed in anion-conducting MerMAIDs, explained by accumulation of a long-lived M intermediate with an unusual short-wavelength maximum absorption peak at 364 nm (7). In both *Ra*CCR1 and *Rs*CCR1 two distinct UV-absorbing intermediates were accumulated upon illumination, one at 390 nm, typical of M intermediates, and the other a triple-peaked species with a uniquely far blue-shifted spectrum with a 330-nm maximum.

Flash photolysis measurements revealed an extremely fast (<1 μs after the flash) appearance of the L intermediate that might be in equilibrium with K during the first 100 μs in *Ra*CCR1 and *Rs*CCR1. The typical M intermediate absorbing at 390 nm was formed during 0.1-1 ms. We could not follow the appearance of the second blue-shifted intermediate (P330) because the low signal-to-noise ratio in the near UV rangelimitedmeasurements to wavelengths ≥ 380 nm. However, we observed bleaching in the entire visible range on the millisecond time scale, which likely indicated accumulation of P330. A decrease of absorption in the blue range was observed during the fast recovery of the unphotolyzed states of both *Ra*CCR1 and *Rs*CCR1. This most probably reflects dissipation of an N-like intermediate that appears earlier in the photocycle.

The short-wavelength absorption of photointermediate P330 is unique among photocycles of microbial rhodopsins. The extremely short wavelength absorption and very high ethylenic stretch wavenumber of a corresponding band in Raman spectra suggest an extremely hydrophobic environment of the retinal moiety. The chromophore in P330 could be, for example, retro-retinal, a derivative in which all double bonds are shifted towards the ring by one position (22), or a free retinal that remains in the binding pocket (23). A linear photocycle involving P330 is the simplest scheme that fits our results; however, we cannot exclude a branched photocycle as was proposed for the *Cr*ChR2_C128T mutant (22).

Rapid desensitization observed in *Ra*CCR1 and *Rs*CCR1 under continuous illumination would potentially allow temporally precise neuronal activation even in the presence of light that can be used for fluorescent imaging. Additional advantages of these two BCCRs are their relatively red-shifted absorption and high Na^+^/H^+^ permeability ratio. Better understanding of their molecular mechanisms will facilitate their rational design for optogenetic needs.

## MATERIALS AND METHODS

### Bioinformatics

BCCR homologs were identified by searching cryptophyte transcriptomes from the MMETS sequencing project (15) and 1KP project (16) using probabilistic inference methods based on profile hidden Markov models (profile HMMs). Profile HMMs were built from previously known BCCR sequences using HMMER software (version 3.1b2; (14)) with default parameters and refined upon functional testing of the homologs by patch clamping. Search procedures were automated with Python 2.7 and the Biopython module (24). The protein sequence alignment was created using MUSCLE algorithm implemented in DNASTAR Lasergene (Madison, WI) MegAlign Pro software. The phylogenetic tree was visualized using Dendroscope software (25).

### Molecular biology

For expression in HEK293 cells, DNA polynucleotides encoding the BCCR opsin domains optimized for human codon usage were synthesized (GenScript, Piscataway, NJ) and cloned into the mammalian expression vector pcDNA3.1 (Life Technologies, Grand Island, NY) in frame with an EYFP tag. For expression in *Pichia*, the opsin-encoding constructs were fused in frame with a C-terminal eight-His tag and subcloned into the pPIC9K vector (Invitrogen). Mutants were generated with Quikchange XL kit (Agilent Technologies, Santa Clara, CA) and verified by sequencing.

### HEK293 transfection and patch clamp recording

HEK293 cells were transfected using the ScreenFectA transfection reagent (Waco Chemicals USA, Richmond, VA). All-*trans*-retinal (Sigma) was added at the final concentration of 3 μM immediately after transfection. Photocurrents were recorded 48-96 h after transfection in the whole-cell voltage clamp mode with an Axopatch 200B amplifier (Molecular Devices, Union City, CA) using the 10 kHz low-pass Bessel filter. The signals were digitized with a Digidata 1440A using pClamp 10 software (both from Molecular Devices). Patch pipettes with resistances of 2-4 MΩ were fabricated from borosilicate glass. The standard pipette solution contained (in mM): KCl 126, MgCl_2_ 2, CaCl_2_ 0.5, Na-EGTA 5, HEPES 25, pH 7.4. The standard bath solution contained (in mM): NaCl 150, CaCl_2_ 1.8, MgCl_2_ 1, glucose 5, HEPES 10, pH 7.4. A 4 M KCl bridge was used in all experiments, and possible diffusion of Cl^−^ from the bridge to the bath was minimized by frequent replacement of the bath solution with fresh buffer. For measurements of the reversal potential shifts under varied ionic conditions, Na^+^ was substituted for K^+^ in the pipette solution to minimize the number of ionic species in the system. To reduce the Cl^−^ concentration in the bath, NaCl was replaced with Na-aspartate; to reduce the Na^+^ concentration, with N-methyl-D-glucamine chloride; to increase the H^+^ concentration, pH was adjusted with H_2_SO_4_. The holding voltages were corrected for liquid junction potentials calculated using the Clampex built-in LJP calculator (26). Continuous light pulses were provided by a Polychrome V light source (T.I.L.L. Photonics GMBH, Grafelfing, Germany) in combination with a mechanical shutter (Uniblitz Model LS6, Vincent Associates, Rochester, NY; half-opening time 0.5 ms). The maximal quantum density at the focal plane of the 40× objective was 7.7 mW mm^−2^ at 515 nm. The action spectra were constructed by calculation of the initial slope of photocurrent and corrected for the quantum density measured at each wavelength. Laser excitation was provided by a Minilite Nd:YAG laser (532 nm, pulsewidth 6 ns, energy 12 mJ; Continuum, San Jose, CA). The current traces were logarithmically filtered using a custom software, and the laser artifact was digitally subtracted. Curve fitting was performed by Origin Pro software (OriginLab Corporation, Northampton, MA).

### Expression and purification of BCCRs from Pichia

The plasmids encoding BCCRs were linearized with SalI and used to transform *P. pastoris* strain SMD1168 (*his4, pep4*) by electroporation. Transformants were first screened for their ability to grow on histidine-deficient medium, and second, for their geneticin resistance. Single colonies that grew on 4 mg/ml geneticin were screened by small-scale cultivation, and clones of the brightest color were selected. For protein purification, a starter culture was inoculated into buffered complex glycerol medium until A600 reached 4–8, after which the cells were harvested by centrifugation at 5000 rpm and transferred to buffered complex methanol medium supplemented with 5 μM all-*trans* retinal (Sigma Aldrich). Expression was induced by the addition of 0.5% methanol. After 24-30 h, the cells were harvested and disrupted in a bead beater (BioSpec Products, Bartlesville, OK) in buffer A (20 mM sodium phosphate, pH 7.4, 100 mM NaCl, 1 mM EDTA, 5% glycerol). After removing cell debris by low-speed centrifugation, membrane fragments were collected by ultracentrifugation at 40,000 rpm in a Ti45 rotor, resuspended in buffer B (20 mM Hepes, pH 7.4, 300 mM NaCl, 5% glycerol) and solubilized by incubation with 1.5% dodecyl maltoside (DDM) for 1.5 h or overnight at 4°C. Non-solubilized material was removed by ultracentrifugation at 50,000 rpm in a TLA-100 rotor. The supernatant was mixed with nickel-nitrilotriacetic acid agarose beads (Qiagen, Hilden, Germany) and loaded on a column. The proteins were eluted with buffer C (20 mM Hepes, pH 7.4, 300 mM NaCl, 5% glycerol, 0.02% DDM) containing 300 mM imidazole. The pigments were concentrated and imidazole was removed by repetitive washing with imidazole-free buffer C using YM-10 centrifugal filters (Amicon, Billerica, MA).

### Absorption spectroscopy and flash photolysis

Absorption spectra of purified BCCRs were recorded using a Cary 4000 spectrophotometer (Varian, Palo Alto, CA). The pK_a_ was determined by fitting the classical Henderson-Hasselbalch equation in the form y = A/(1+10E(pK_a_−pH)) to experimental data. Light-induced absorption changes were measured with a laboratory-constructed crossbeam apparatus. Excitation flashes (532 nm, 6 ns, 40 mJ) were provided by a Surelite I Nd-YAG laser (Continuum, Santa Clara, CA). Measuring light was from a 250-W incandescent tungsten lamp combined with a McPherson monochromator (model 272, Acton, MA). Absorption changes were detected with a Hamamatsu Photonics (Bridgewater, NJ) photomultiplier tube (model R928), protected from excitation laser flashes by a second monochromator of the same type. Signals were amplified by a low noise current amplifier (model SR445A; Stanford Research Systems, Sunnyvale, CA) and digitized with a GaGe Octopus digitizer board (model CS8327, DynamicSignals LLC, Lockport, IL), maximum sampling rate 50 MHz. Logarithmic filtration of the data was performed using the GageCon program (27).

### Fourier-transformed Raman spectroscopy

Fourier-transformed Raman spectra were collected in 5 μl of a concentrated detergent-solubilized protein in pH-adjusted elution buffer placed in a metallic holder and covered by adhesive tape. The scattering was recorded in 180° backscattering geometry, using FRA106/s accessory to the Bruker IFS66vs spectrometer, with Nd-YAG laser excitation provided at 1064 nm, at a 2 cm^−1^ resolution, controlled by the OPUS software. At least 10000 scans averaged per sample. Raman spectra of the buffers were taken separately in the same geometry and subtracted to get pure protein spectra.

### Statistics

Descriptive statistics was used as implemented in Origin software. The data are presented as mean ± s.e.m. values; the data from individual replicates are also shown when appropriate. The sample size was estimated from previous experience and published work on similar subjects, as recommended by the NIH guidelines (28).

### Data availability

The polynucleotide sequences of BCCR homologs reported in this study have been deposited to GenBank (accession numbers MN585290-MN585308).

## Acknowledgements

We thank Olivier Morelle for technical assistance with bioinformatics analysis. This work was supported by National Institutes of Health Grant R01GM027750, the Hermann Eye Fund, and Endowed Chair AU-0009 from the Robert A. Welch Foundation to J.L.S, and by the Natural Sciences and Engineering Research Council of Canada (NSERC) Discovery Grant RGPIN-2018-04397 to L.S.B. The content is solely the responsibility of the authors and does not necessarily represent the official views of the National Institutes of Health.

## SUPPLEMENTARY FIGURE LEGENDS

**Figure S1.** Protein sequence alignment of the opsin domains of BCCRs first reported in this study. Residues are color-coded according to their chemical properties.

**Figure S2.** (A) Amplitudes of BCCR photocurrents recorded at −60 mV at the amplifier output upon photostimulation at the peak sensitivity wavelength. Stationary currents were measured at the end of a 1-s light pulse. The data are the mean values ± sem recorded from 3-24 individual cells. The data from individual cells are shown as empty diamonds. (B) The action spectra of photocurrents. The data are the mean values ± sem recorded from 6-8 individual scans.

**Figure S3.** Representative photocurrent traces recorded from indicated BCCRs at −60 mV at the amplifier output in response to a light pulse, the duration of which is shown as a colored bar on top.

**Figure S4.** (A-C) The reversal potentials measured under indicated ionic conditions. The data points are the mean values ± sem. The data from individual cells are shown as empty diamonds.

**Figure S5.** (A and B) The difference spectra (treated minus untreated sample) obtained by incubation of indicated proteins with hydroxylamine (HA) in the dark (black lines) and under illumination (red lines). (C and D) The time course of hydroxylamine bleaching of the UV and visible absorption bands. The bar on top shows the time of illumination.

**Figure S6.** (A) The rate of absorbance recovery at 530 nm of *Ra*CCR1 in *Pichia* membranes (red line) as compared to that in detergent (black line). (B) The UV portion of the *Rs*CCR1 difference spectrum (that of *Ra*CCR1 from Fig. 1B in the main text is shown for comparison). (C) Peak amplitude of *Ra*CCR1 photocurrents in response to 0.1 Hz trains of 6-ns laser flashes.

**Figure S7.** (A) A series of *Ra*CCR1 current traces in response to 1-s light pulses of different intensities. (B and C) The dependence of peak current amplitude and desensitization on the stimulus intensity in *Ra*CCR1 (B) and *Rs*CCR1 (C).

**Figure S8.** (A) A series of flash-induced absorption changes recorded at the indicated wavelengths between 390 and 570 nm in purified *Rs*CCR1. (B) Absorption changes at the wavelengths corresponding to the K and L intermediates in the microsecond range in *Rs*CCR1.

**Figure S9.** (A) Mean photoinduced absorbance changes recorded from purified *Ra*CCR1 in the 0-100 μs time window. (B-F) Spectral transitions in *Ra*CCR1 derived by global fit analysis.

**Figure S10.** A laser-flash-evoked photocurrent trace recorded from *Ra*CCR1 (the black dashed line) at the holding potential near the equilibrium potential for Na^+^. The red solid line shows a multiexponential fit.

